# Spatial cueing effects are not what we thought: on the timing of attentional deployment

**DOI:** 10.1101/2020.04.08.031302

**Authors:** Itay Yaron, Dominique Lamy

## Abstract

Extensive research has shown that objects that are salient or match our task goals are most likely to capture attention. But are we at the mercy of the constant changes occurring in our environment, and automatically move our attention to the ever-changing location with the highest priority? Or do we wait for clues that the appropriate moment has arrived to deploy our attention? We addressed this hitherto neglected issue in three experiments. Using a spatial-cueing paradigm, we examined whether attention is deployed as soon as a salient change occurs (the cue), or only when the context signaling that attention should be deployed appears (the search display). The cue matched the target color and was therefore expected to enjoy high attentional priority. We used two separate response compatibility manipulations, one pertaining to the cue, in the cueing display, and the other to the cued distractor, in the search display. Neutral conditions allowed us to disentangle the respective effects of these manipulations. Our results support the hypothesis that attention does not occur until the search-relevant context appears. These findings challenge the traditional interpretation of spatial-cueing effects. They are discussed within the Priority Accumulation Framework (PAF) that we confront to other attention models.

---

We need attention to recognize objects but can attend to very few objects at any given time. To explain why we are usually fast at finding potentially important objects in our environment despite such limitations of our perceptual system, most models suggest that powerful mechanisms, driven by stimulus salience and the observers’ goals, guide our attention (e.g., Cave & Wolfe, 1990; Folk, Remington, & Johnston, 1992; Itti & Koch, 2000; Theeuwes, 2010; Wolfe & Horowitz, 2017; Yantis & Egeth, 1999). These mechanisms operate in parallel during an initial capacity-unlimited stage often referred to as the preattentive stage (Neisser, 1976), and determine the overall priority level of each location. Then, during a second, capacity-limited stage, attention is deployed to the highest-priority location.

There has been intense research to assess what factors determine the distribution of attentional priorities across the visual field (for recent reviews, see e.g. Lamy, Leber, & Egeth, 2012; Theeuwes, 2010; Wolfe & Horowitz, 2017). By contrast, researchers have remained largely silent on the critical issue of how the capacity-limited stage of attentional deployment is triggered. Are we at the mercy of the constant changes occurring in our environment, and automatically move our attention to the ever-changing location with the highest priority? Or do we wait for clues that the appropriate moment has arrived to expend our scarce attentional resources? The latter behavior would shield our attentional system against relentlessly shifting our processing resources to potentially irrelevant events. Yet, most researchers implicitly assume the former behavior to be the rule.

For instance, consider the standard interpretation of the spatial cueing paradigm, which is widely used to investigate what objects capture our attention against our will. On a typical trial, observers look for a target defined by a given property (e.g., its color). Shortly before the search display is presented, a cue (e.g., an abruptly onset object) appears at one of the potential target locations. Finding faster search performance when the target appears at the same location as the cue (valid-cue trials) than at a different location (invalid-cue trials), is taken to indicate that the cue captured attention: because the cue is the highest-priority object when the cueing display appears, attention is automatically deployed to its location (e.g., Folk et al., 1992; Folk & Remington, 2006; Gaspelin, Ruthruff, & Lien, 2016; Theeuwes, Atchley, & Kramer, 2000).

Yet, is it not surprising that participants should fail to use their knowledge of the regularities inherent to the task? The cueing and search displays have consistent and easily distinguishable temporal and visual characteristics. Previous research has shown that participants can use temporal regularities to deploy their attention at the most appropriate time (e.g., Coull & Nobre, 1998; Lamy, 2005) and are sensitive to display-wide characteristics (Gibson & Kelsey, 1998). Thus, in a typical spatial cueing paradigm, after just a few trials, participants should learn that the target always appears in the search display and never in the cueing display. Accordingly, they should deploy their attention only after detecting the context most likely to signal the target presence, that is, the search display.

Recently, we suggested a new model of attentional allocation, the Priority Accumulation Framework (PAF), which stipulates that the attentional-deployment stage is not inflexibly initiated in response to moment-to-moment changes in attentional priorities, but is instead triggered by contextual information (Gabbay, Zivony, & Lamy, 2019; Lamy, Darnell, Levi & Bublil, 2018). We proposed this model in order to accommodate findings that could not be explained within the framework of the standard interpretation of spatial cueing effects (see Gabbay et al., 2019; Lamy et al., 2018 for details).

According to PAF, the attentional priority accruing to a given location mainly depends on how similar the successive objects that have appeared at that location (e.g., a cue, a distractor or the target) are to the target, and on the physical salience of these objects. Priority weights accumulate over time at each location, until the search context (i.e., the search display) signals that selection can occur. The first attentional shift is then made to the item that wins the competition (i.e., the item with the highest accumulated attentional priority). How long it takes for the competition to be resolved varies as a function of how large the winner’s leading edge is.

An important implication of this scheme is that one may observe cue validity effects even if attention was never directed to the cue. Specifically, the target may have the most highly activated location even when it is not cued, and in that case, it is the object that receives the first shift of attention. However, the resolution of the competition that leads to its selection is nevertheless faster if the target (valid-cue trials) rather than a distractor (invalid-cue trials) benefits from the extra activation provided by the cue – hence the cue validity effect. Thus, cue validity effects do not necessarily index attentional deployment. By contrast, the compatibility between the response associated with the cued distractor and the response associated with the target attests that enhanced processing took place at the cued location and therefore, that this location was selected. Thus, response compatibility effects are a reliable index of attentional deployment.

The objective of the present study was to test PAF’s hypothesis that attention is not directed to the object with the highest priority at any given time, but only when contextual information signals that selection should occur. In operational terms, we asked whether in a spatial cueing search task, attentional deployment, indexed by response compatibility effects, occurs in the cue display (as the standard interpretation of spatial cueing effects would predict) or in the search display (as PAF would predict).

Each search display contained four arrows, each in a different color and pointing in a different direction (see Figure 1). The target was defined by a specific color (e.g., red) and participants had to respond to its direction (e.g., left or right). Thus, one distractor was associated with the response opposite to the target (incompatible distractor) and the other two were associated with no response (neutral distractors). The critical aspect of the present study is that the cue also could be associated with a task-relevant response. The cueing display included a set of four small, identically oriented arrows surrounding each potential target location. One of them (the cue) was in the target color and the other were white. The cue arrows could point in any of the four possible directions. Thus, the cue arrows could be compatible, incompatible or neutral relative to the target arrow direction. The other three arrow groups pointed in neutral directions.

**Figure 1:**
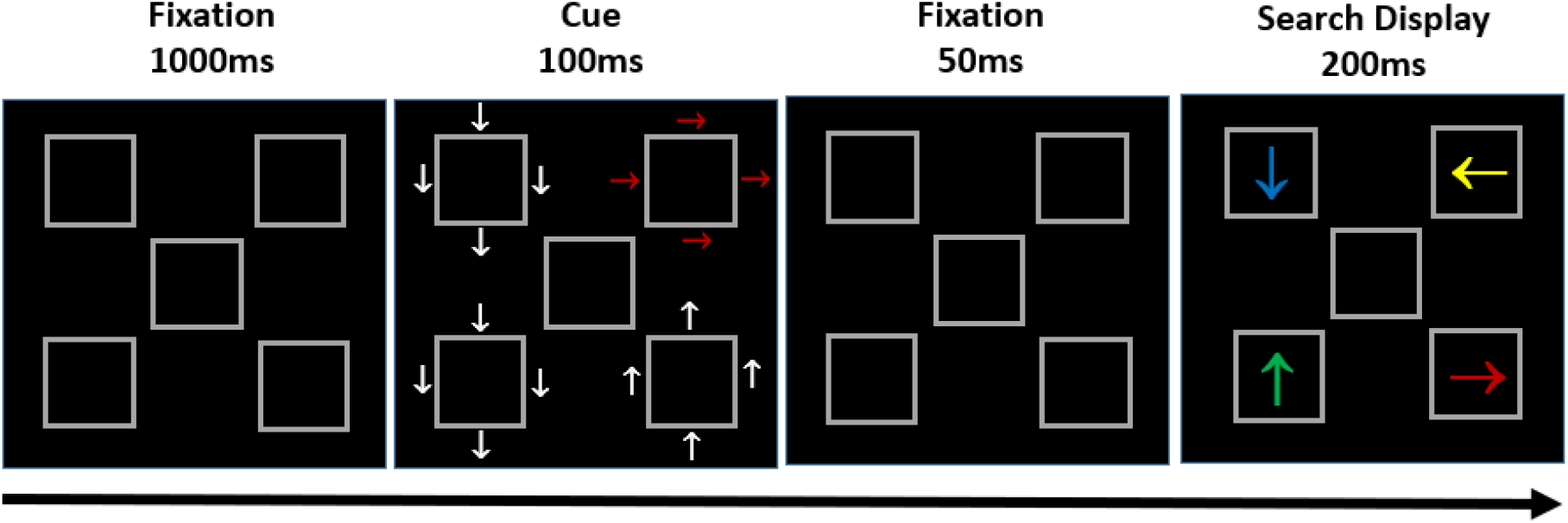
Sample sequence of events, for the red-horizontal target group. Participants searched for the red arrow in the search display and reported its orientation (left or right). This example corresponds to an invalid-cue trial, in which the cue was compatible with the target and the cued item was an incompatible distractor.

Because the cue always shared the target color, we assumed that the cue location had the highest priority and therefore, we expected reliable cue validity effects - in line with the contingent capture hypothesis (e.g., Folk et al., 1992; Folk & Remington, 1998). The question of interest was whether attention would be deployed to the cue location in the cueing display or in the search display.

We used response compatibility effects to measure attentional deployment. We relied on the premise that the location to which attention is deployed enjoys enhanced processing (e.g., Kahneman & Henik, 1981; Treisman & Gelade, 1980). Thus, if attention is deployed in the cue display, participants should process the direction of the cue, which should therefore affect responses to the target arrow (that is, induce response compatibility effects): responses to the target should be faster when the cue arrows point in the same direction as the target than when they point in the alternative direction. Likewise, if attention is deployed in the search display, the direction of the distractor that appeared at the location of the cue (cued distractor) should affect responses to the target arrow: responses to the target should be faster when the distractor that appears at the cued location points in a neutral direction than when it points in the direction opposite to the target’s.

Several authors already reported response compatibility effects associated with the distractor at the (invalidly) cued location in the search display (e.g., Carmel & Lamy, 2014; Theeuwes et al., 2000; Zivony & Lamy, 2018). However, these authors construed such compatibility effects within the standard interpretation of the spatial cueing paradigm. They assumed – yet did not demonstrate - that attention was deployed to the cue in the cueing display and still dwelled at the cue location when the search display came on; hence, the features of the cued distractor were processed, accounting for the response compatibility effect.

The design of the present study allowed us to directly test this hypothesis against PAF’s by measuring the critical effects (cue compatibility – in the cue display, and cued distractor compatibility – in the search display) separately. Note, however, that these effects could interact and render the interpretation of the results unwieldy. As is explained next, in the statistical analyses section, we performed a set of analyses designed to overcome this potential problem.

## Statistical Analyses

In all three experiments of the present study we conducted two sets of analyses.

### Planned comparisons

We first conducted planned comparisons to assess the two most critical effects in this study, cue compatibility and cued-distractor compatibility when they were not contaminated by each other. Specifically, we measured (1) the cue compatibility effect (compatible vs. incompatible) when the cued item (in the search display) was a neutral distractor and (2) the cued-distractor compatibility effect (incompatible vs. neutral) when the cue (in the cueing display) was neutral. To estimate the strength of the evidence in favor or against the null hypothesis for these two effects, we performed Bayesian analyses, using the anovaBF function from the BayesFactor package in R (Morey, Rouder, & Jamil, 2015), with the r = .707 prior. Following Dienes and Mclatchie, (2018), we consider BF_10_ to provide conclusive evidence against the null hypothesis if it is larger than 3, in favor of the null if it is smaller than .33, and to be inconclusive if it stands between .33 and 3.

### Overall ANOVA

For completeness, we also conducted an analysis of variance (ANOVA) with cue compatibility (compatible, neutral and incompatible) and cued item in the search display (target, neutral distractor and incompatible distractor) as within-subject factors. The effect of the cued item (i.e., whether the location of the cue coincided with the location of the target, a neutral distractor or the incompatible distractor in the search display) includes two components: the cue-validity effect that is not contaminated by response compatibility (target-cued vs. neutral-distractor cued trials) and the cued-distractor compatibility effect (neutral- vs. incompatible-distractor cued trials)^1^. We reported separate analyses of these two components.

In order to clarify any interaction between cue compatibility and cued item compatibility, we conducted a separate analysis of the cue compatibility effect (compatible, neutral, incompatible) when the cued item was the target and when it was an incompatible distractor. The cue compatibility effect when the cued item was neutral was reported earlier, in the planned comparisons. Whenever the cue compatibility effect was significant, we also reported the contrast of main interest, namely, the contrast between compatible- and incompatible-cue trials.

## Experiment 1

### Methods

#### Participants

##### Sample-size selection

We relied on the size of effect of the response compatibility between the target and the cued distractor in Carmel and Lamy’s study (2014, Experiment 2, target-color cue), which is similar to the cued item compatibility effect in the present study. We conducted this analysis with G*Power (Faul, Erdfelder, Buchner, & Lang, 2013) using an alpha of .05, power of .95, and the effect size reported by Carmel and Lamy 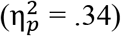. We found the minimum required sample size to be 9 participants. Thus, we were confident that a sample of 14 participants would provide enough power. We used the same number of participants in all three experiments.

Fourteen Tel Aviv University students (12 females, mean age = 26 years, SD = 4.3) volunteered to participate in Experiment 1. All participants were right-handed and reported normal or corrected-to-normal visual acuity. All protocols for this and the following experiments were approved by the Tel Aviv University ethics committee.

#### Apparatus

The experiment took place in a dimly lit room. Stimuli were presented on 23’ screen, using 1920×1080 resolution graphics mode and 120-Hz refresh rate. Responses were collected via the computer keyboard. Viewing distance was set at approximately 60 cm from the monitor.

#### Stimuli

The sequence of events is presented in Figure 1. The fixation display consisted of five gray square outline placeholders (2.4° × 2.4°), one at the center of the screen (fixation) and the remaining four equally spaced at the corners of an imaginary square (7.2°×7.2°). Thus, the distance between the central- and outer-frame centers distance was 3.4°. The cue and search displays were similar to the fixation display, except for the following changes.

In the cue display, a group of four small arrows (.8° × .8°) pointing in the same direction (left, right, up or down) was added around each of the four outer placeholders, in a diamond configuration (3° × 3°). One group of arrows (the cue) was red for half of the participants and green for the other half and could point in any of the four possible directions. The remaining three groups of arrows were white and pointed in two directions orthogonal to the possible target directions (i.e., up or down for participants assigned to horizontal targets and left or right for participants assigned to vertical targets – see description of the search display below for further explanations), with two white groups pointing in one direction and the remaining white group in the other direction.

In the search display, an arrow (2°×1.2°), appeared in the center of each outer placeholder and each pointed in a different direction (left, right, up and down). The target was defined by its color, which was always in the same as the cue color (i.e., red for half of the participants and green for the other half). The remaining three arrows (the distractors) were in different colors (blue, orange and either red or green for the green- and red-target participants, respectively). For half of the participants the direction of the target arrow was always horizontal (left or right) and for the other half, it was always vertical (up or down). Color RGB coordinates were (128, 128, 128) for gray, (255, 0, 0) for red, (0, 255, 0) for green, (0,123,255) for blue and (222,125,0) for orange, and (255,255,255) for white.

#### Procedure

Participants had to search for the color-defined target arrow in the search display and report its orientation. They were asked to respond as fast and as accurately as possible by pressing the key ‘M’, or the key ‘O’ with their right hands on the computer keyboard when the target pointed to the left or to the right, respectively, in the horizontal-target group, and when it pointed downwards or upwards, respectively, in the vertical-target group.

Each trial began with the fixation display for 1,000ms, followed by the cue display for 100ms and then again, by the fixation display for 50ms. Then, the search display appeared for 200ms and was followed by another fixation display, which remained on the screen for 1,800 ms or until response, whichever came first. Following an incorrect response or response timeout, participants heard an error beep (225 Hz) for 300ms.

#### Design

The experiment consisted of 15 practice trials, followed by 6 blocks of 98 trials each. Target color (red or green) and directions (horizontal or vertical) were randomly assigned between participants. Cue and target locations were randomly selected and were therefore uncorrelated. In the cueing display, the cue arrows direction was compatible with the target direction on 1/3 of the trials, incompatible with it on 1/3 of the trials and neutral on 1/3 of the trials (1/6 for each of the two possible neutral directions). In the search display, the target arrow was equally likely to point to the right or left, for the horizontal group and downward or upwards, for the vertical-target group. The remaining possible arrow directions in the search display were randomly assigned to the remaining locations. Thus, one distractor arrow’s direction was incompatible with the target direction, while the remaining two distractors were neutral.

### Results

No participant met the conditions for exclusion (either an average reaction time or an average accuracy rate differing from the group’s mean by more than 3 standard deviations). Error trials (5.05% of the trials) as well as RT outliers, defined as any correct-response trial with an RT differing from the mean of its cell by more than 2.5 standard deviations (2.05 % of the remaining trials) were excluded from all RT analyses. Mean RTs and accuracy data are presented in Table 1.

**Table 1.** Mean reaction times (in milliseconds) and mean accuracy (in percentage) in Experiment 1, as a function of the compatibility of the cue with the target (Cue Compatibility: compatible, incompatible, neutral) and of the cued item in the search display (Cued Item: target, incompatible distractor, neutral distractor).

|  | Cued Item |  |  |
| --- | --- | --- | --- |
|  | *Target* | *Neutral distractor* | *Incompatible distractor* |
| Cue Compatibility |  |  |  |
| *Reaction times* |  |  |  |
| Compatible Cue | 479.2 (17.4) | 548.8(10.3) | 569.2 (13.1) |
| Neutral Cue | 492.3(6.5) | 541.5 (7.5) | 561.8(8.4) |
| Incompatible Cue | 512.2(11.6) | 546.1(9.7) | 557.9(12.5) |
| *Accuracy* |  |  |  |
| Compatible Cue | 98.1 % (2.9) | 95.2 % (1.8) | 89.0 % (4.7) |
| Neutral Cue | 97.5 % (2.15) | 96.6% (1.0) | 91.9% (2.4) |
| Incompatible Cue | 97.0 % (1.7) | 94.9 % (1.7) | 92.6 % (2.6) |

*Note:* Within-subject standard errors (Morey, 2008) are presented in parentheses.

#### Planned comparisons

##### Reaction times

When the cue (in the cueing display) was neutral, the effect of cued distractor compatibility (i.e., incompatible vs. neutral distractor at the cue location in the search display) was highly significant, F(1, 13) = 14.58, p = .002, 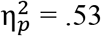. A Bayesian analysis confirmed that the evidence for this effect was conclusive, *BF*_10_ = 14.41 (see Figure 2). When the cued item (in the search display) was a neutral distractor, the effect of cue compatibility (i.e., compatible vs. incompatible cue in the cueing display) was not significant, F < 1. A Bayesian analysis confirmed that the evidence for this null effect was conclusive, *BF*_10_ = .27 (see Figure 3).

**Figure 2:**
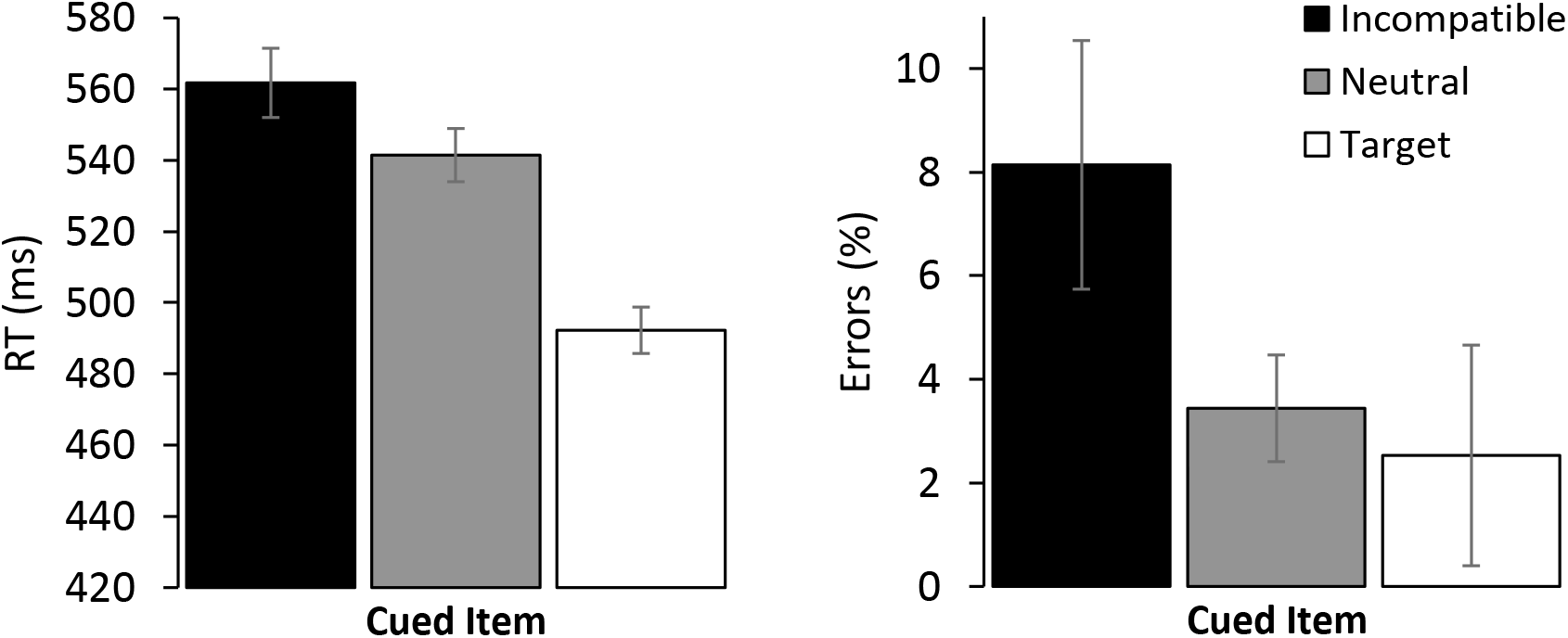
Mean reaction times (RT - left panel) and mean errors (right panel) in Experiment 1 for trials in which the cue compatibility was neutral, as a function of the cued item. Error bars denote Within-subject standard errors (Morey, 2008).

**Figure 3:**
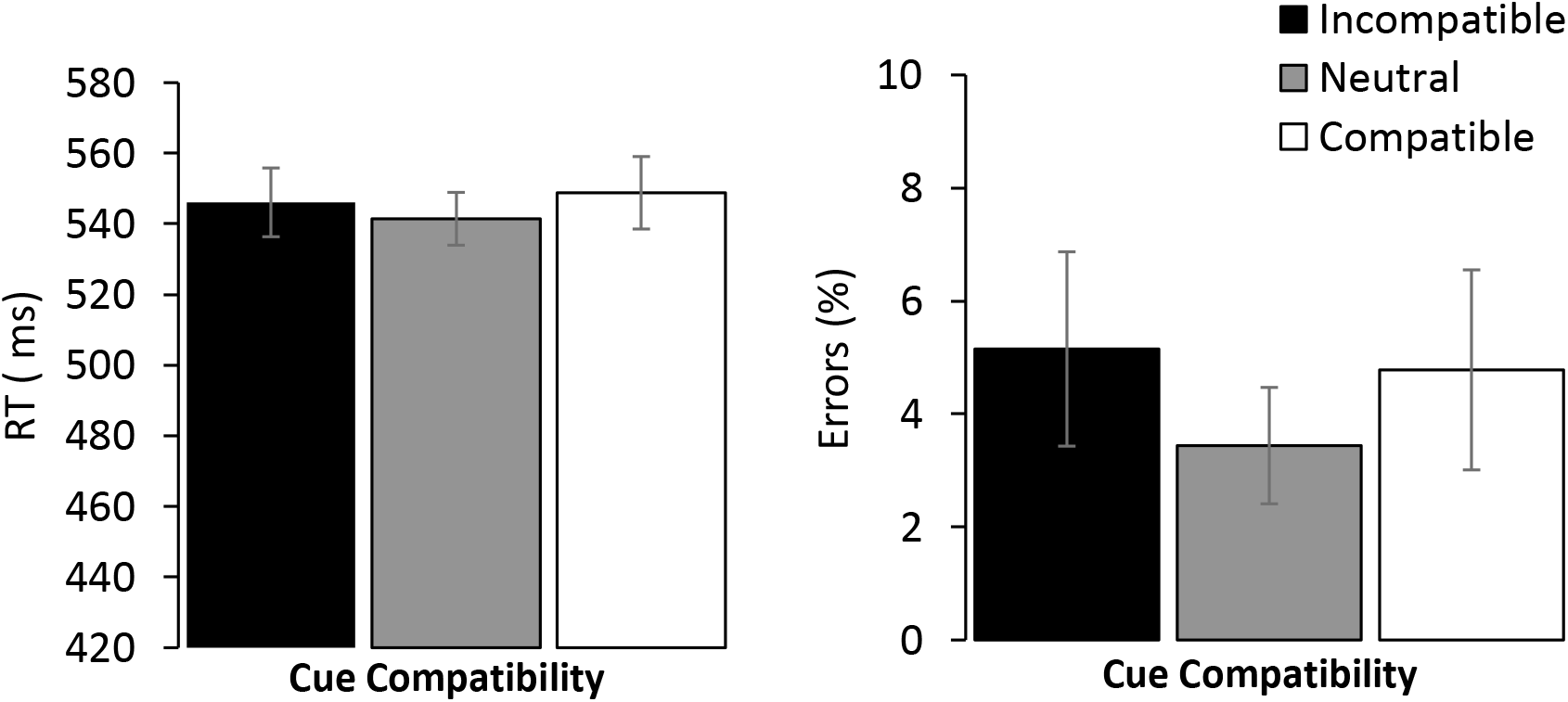
Mean reaction times (RT - left panel) and mean errors (right panel) in Experiment 1 for trials in which the cued item was neutral, as a function of cue compatibility. Error bars denote Within-subject standard errors (Morey, 2008).

##### Accuracy

The accuracy data closely mirrored the RT data. When the cue was neutral, the effect of cued distractor compatibility was significant, F(1, 13) = 16.76, p =.001, 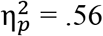, with conclusive evidence for this effect, *BF*_10_ = 26.57. When the cued item was a neutral distractor, the effect of cue compatibility was not significant, F(2, 26) = 1.35, p =.2, 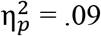, with conclusive for the null, *BF*_10_ = .27.

#### Overall ANOVA

##### Reaction times

The main effect of cued item (which item in the search display appeared at the location of the cue) was highly significant, F(2, 26) = 203.7, p < .0001, 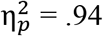: RTs were faster when the target rather than a neutral distractor appeared at the cued location, F(1, 13) = 241.91, p < .0001, 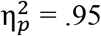 (cue-validity effect) and when a neutral rather than an incompatible distractor appeared there, F(1, 13) = 50.15, p < .0001, 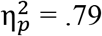 (cued-distractor compatibility effect). There was no main effect of cue compatibility, F < 1. The interaction between cue compatibility and cued item was significant, F(4, 52) = 14.08, p < .0001, 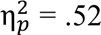.

Further analyses clarified this interaction. When the cued item was the target, the cue compatibility effect was significant, F(2, 26) = 7.59, p = .003, 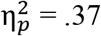. A paired comparison revealed that RTs were slower when the cue was incompatible than when it was compatible, F(1, 13) = 7.97, p = .014, 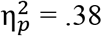. When the cued item was the incompatible distractor, the cue compatibility effect was not significant, F(2, 26) = 1.05, p = .36, 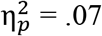.

##### Accuracy

The numerical trends on accuracy were similar to those observed on the RT data, thus removing any concern for a speed-accuracy trade-off. The main effect of cued item was significant, F(2, 26) = 12.97, p < .0001, 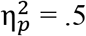, with significant effects of both its cue-validity and cued-distractor compatibility components, F(2, 26) = 5.86, p =.03, 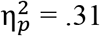 and F(2, 26) = 17.14, p = .001, 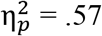, respectively. Neither the main effect of cue compatibility nor the interaction between the two factors was significant, F < 1 and F(4, 52) = 2.13, p = .09, 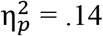, respectively.

### Discussion

Experiment 1 yielded two findings that are predicted both by traditional accounts and by PAF. First, the cue produced a large validity effect. Second, compatibility between the directions of the target and cued distractor arrows (cued-item compatibility, in the search display) strongly modulated performance. These findings indicate that the information presented at the cued location in the search display enjoyed enhanced processing. They are predicted by traditional models such as the contingent capture (Folk et al., 1992), rapid disengagement (Theeuwes, 2010), and attention dwelling (Gaspelin et al., 2016) models. These all posit that attention is deployed to the cue in the cueing display and dwells on its location until the search display appears. These findings are also predicted by PAF (Lamy et al., 2018), which stipulates that attention is deployed after the search display appears, to the location that has accumulated the highest level of priority across the attentional episode Here, that location was most likely to be that of the cued item, because the cue shared the target color.

However, the critical effects, which concern the compatibility between the cue and target, were ambiguous. On the one hand, in line with PAF’s prediction, the cue’s direction had no impact on responses to the target when the cued item was neutral. On the other hand, when the cued item was the target, performance was clearly better on compatible-than on incompatible-cue trials, as predicted by traditional models. Finally, the results observed when the cued object was an incompatible distractor supported neither account: there was a numerical trend towards a reverse effect of the cue-target compatibility, on both RTs and accuracy.

The notion of object-file (Kahneman, Treisman, & Gibbs, 1992; see also Gordon & Irwin, 1996; Mitroff, Scholl, & Noles, 2007; Noles, Scholl, & Mitroff, 2005) or event-file (van Dam & Hommel 2010; Hommel, 1998) may help us resolve the apparent inconsistencies in these cue compatibility effects. An object file is an episodic representation that stores and updates information about a given object over time. When attention is focused on a location, the information that recently appeared at that location and was stored in the corresponding object-file, is automatically retrieved. Identification is faster when the current and previous features of the object cohere than when they mismatch. Although there is some variance in how different authors account for this finding (e.g., Kahneman et al., 1992; Neill, 1997; van Dam & Hommel, 2010), they all refer to a backward process: focusing attention on a given location now, triggers the retrieval of information that was presented at that location earlier.

This episodic memory retrieval account entails that here, the cued object in the search display was identified faster when its direction and the direction of the cue arrows matched than when they mismatched. On valid-cue trials, this explains why performance was better for compatible than for incompatible cues. On trials in which an incompatible distractor was cued, this explains why performance tended to be better when the cue was also incompatible (the successive arrows at the cue location shared the same direction) than when it was compatible (the successive arrows at the cue location had opposite directions). On trials in which a neutral distractor was cued, this explains why performance was similar for compatible- and incompatible cues, which both failed to match the direction of the cued object. In line with this account, performance was best on neutral-cue trials when the cued distractor was also neutral: this trend, although non-significant, was observed on both RTs and accuracy measures. In line with PAF, after the distractor at the cued location was identified (faster or slower, depending on its match with the stimulus at its location in the cueing display), whether it elicited the same response as the target or the opposite response strongly affected performance - hence the cued item compatibility effect observed across conditions.

## Experiment 2

The objective of Experiment 2 was to investigate the aspects of the search context that might trigger the deployment of spatial attention. As is clear from Figure 1, the cue always included a colored object among three white objects, whereas the search display always included four objects of different colors. Thus, the onset of a multi-color display reliably signaled that the search could start. In addition, the displays were always presented in the same temporal sequence: first, the fixation display, then the cue display and finally, the search display, with exactly the same timing. It is well established that temporal information, whether explicit (e.g., Coull & Nobre, 1998) or implicit (e.g., Olson & Chun, 2001) can guide visual attention. Thus, participants could also rely on temporal regularities to identify the search display and deploy their attention.

In Experiment 2, we eliminated these two predictors of the search display onset. Experiment 2 was similar to Experiment 1, except for two changes (see Figure 4). The arrows in the cue display now had the same colors as the arrows in the search display. Thus, while the cue still matched the target-defining feature, it was no longer a singleton. In addition, the search display still followed the cue display as in Experiment 1 on half of the trials, but on the other half, it appeared immediately after the fixation display (i.e., the cue display was omitted and the search display appeared in its stead). We reasoned that these changes should weaken the context cues that signal the appearance of the search display, and that participants might now rely on the detection of the target color (in the cueing display) to deploy their attention. If so, positive cue compatibility effects should emerge across conditions of the cued distractor compatibility, and in particular when the cued item is neutral.

**Figure 4:**
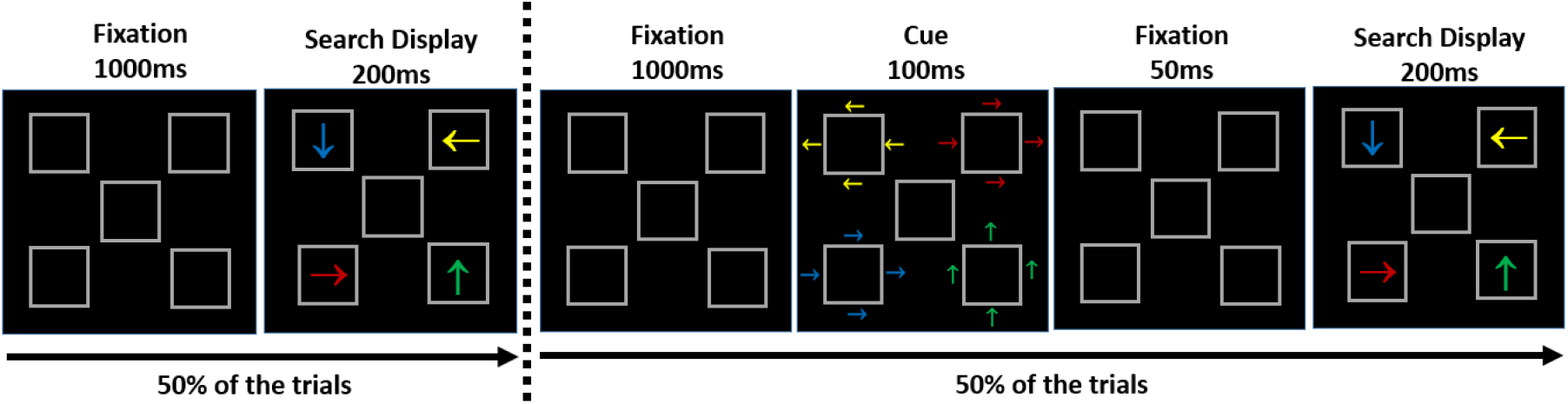
Sample sequence of events in Experiment 2, for the green-vertical target group. Participants searched for the green arrow in the search display and reported its orientation (up or down). *Left panel:* Search-only trial (in which the cue display was omitted). *Right panel:* Regular trial. This example corresponds to a same-location trial in which the cue was compatible with the target response.

### Methods

#### Participants

Fourteen new Tel Aviv University students (8 females, mean age = 28 years, SD = 3.49) volunteered to participate in Experiment 2. All participants except one were right-handed and reported normal or corrected-to-normal visual acuity.

#### Stimuli, procedure, design and statistical analysis

The sequence of events is presented in Figure 4. The stimuli, procedure and design were similar to those of Experiment 1, except for the following changes. On half of the trials, the sequence of events was the same as in Experiment 1 (regular trials), whereas on the other half, the fixation display was directly followed by the search display and the cue display was omitted (search-only trials). Regular and search-only trials were randomly intermixed. In addition, when the cue display was present, one arrow group was in the same color as the target (i.e., green or red, depending on target-color group), while the remaining groups of arrows were in different colors (red/green, respectively, blue and orange). Thus, the cue-display colors were similar to the search-display colors.

### Results

No participant met the conditions for exclusion. Error trials (6.63% of the trials) and RT outliers (2.05 % of the remaining trials) were excluded from all RT analyses. So were search-only trials (in which the cue display was omitted), because these were filler trials that did not allow any informative analyses. Mean RTs and accuracy data are presented in Table 2.

**Table 2.** Mean reaction times (in milliseconds) and mean accuracy (in percentage) in Experiment 2, as a function of the compatibility of the cue with the target (Cue Compatibility: compatible, incompatible, neutral) and of the cued item in the search display (Cued Item: target, incompatible distractor, neutral distractor).

|  | Cued Item |  |  |
| --- | --- | --- | --- |
|  | *Target* | *Neutral distractor* | *Incompatible distractor* |
| *Cue Compatibility* |  |  |  |
| *Reaction times* |  |  |  |
| Compatible Cue | 433.6 (13.1) | 513.2 (10.0) | 544.4 (12.3) |
| Neutral Cue | 459.6 (11.9) | 514.3 (8.0) | 528.1 (7.2) |
| Incompatible Cue | 480.4 (9.5) | 517.9(8.5) | 520.7 (4.2) |
| *Accuracy* |  |  |  |
| Compatible Cue | 96.6 % (2.4) | 94.8 % (1.7) | 89.7 % (3.6) |
| Neutral Cue | 97.2 % (1.9) | 96.3% (1.4) | 89.9% (1.3) |
| Incompatible Cue | 91.2 % (5.4) | 93.5 % (1.6) | 87.1 % (4.1) |

*Note:* Within-subject standard errors (Morey, 2008) are presented in parentheses.

#### Planned comparisons

##### Reaction times

The results of Experiment 2 closely replicated the findings of Experiment 1. When the cue was neutral, the effect of cued distractor compatibility was highly significant, F(1, 13) = 11.19, p = .005, 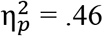, with conclusive evidence for this effect, *BF*_10_ = 7.23 (see Figure 5). When the cued item was a neutral distractor, the effect of cue compatibility was not significant, F < 1, with conclusive evidence for the null, *BF*_10_ = .328 (see Figure 6).

**Figure 5:**
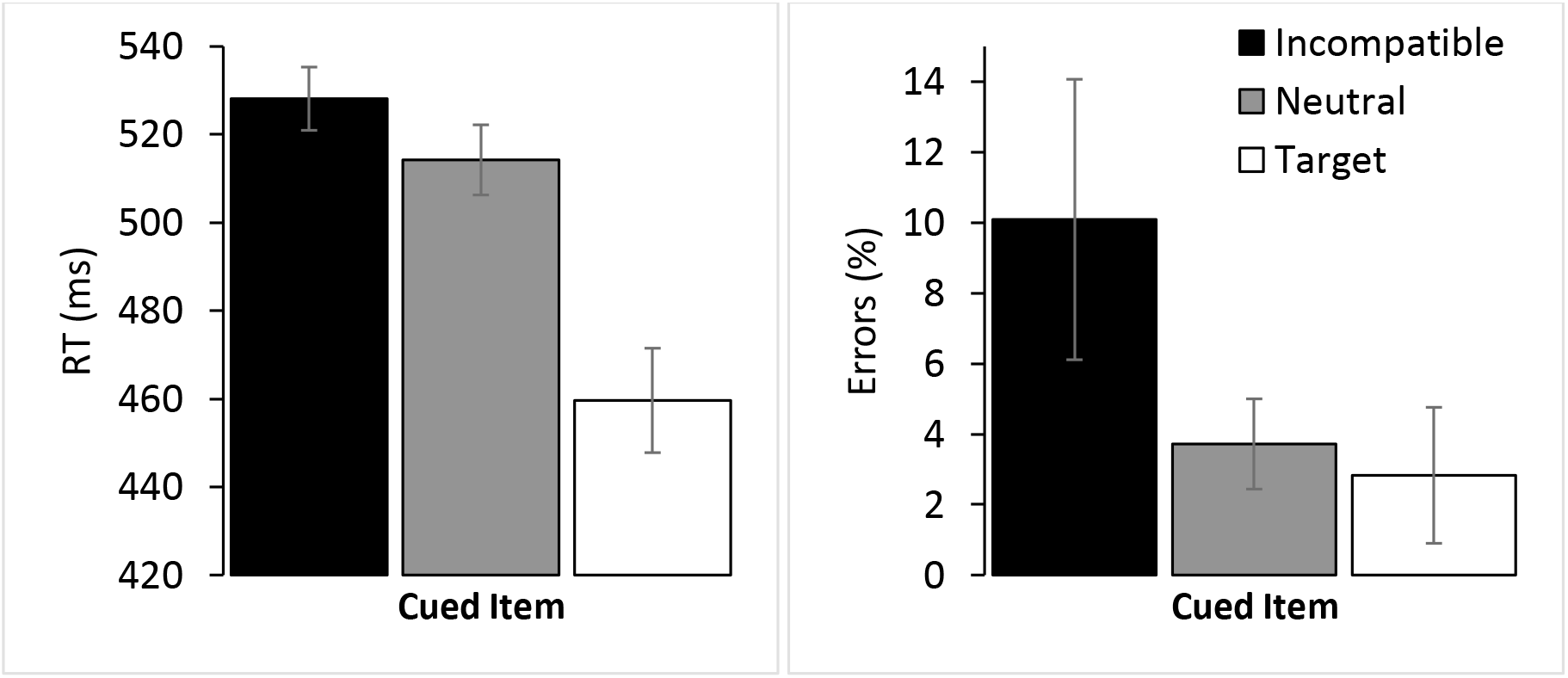
Mean reaction times (RT - left panel) and mean errors (right panel) in Experiment 2 for trials in which the cue compatibility was neutral, as a function of the cued item. Error bars denote within-subject standard errors (Morey, 2008).

**Figure 6:**
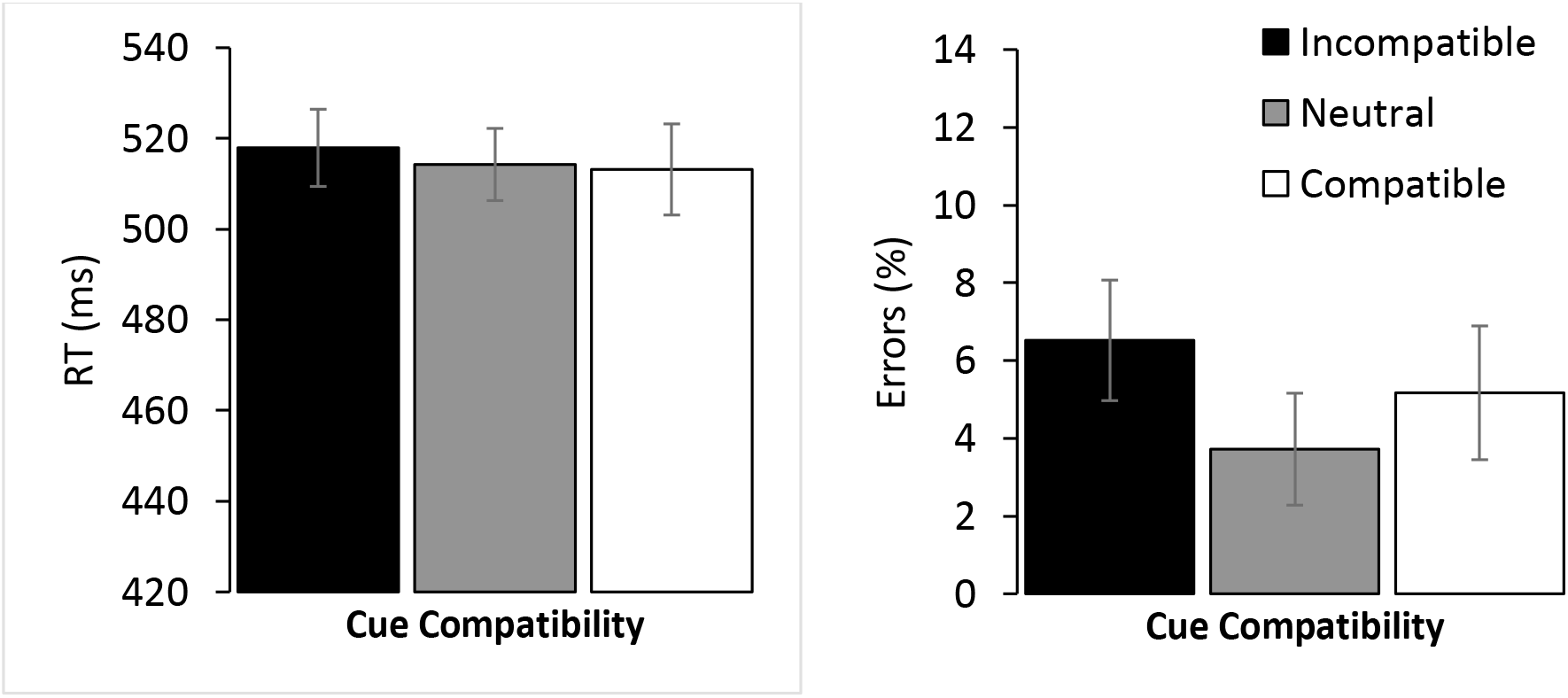
Mean reaction times (RT - left panel) and mean errors (right panel) in Experiment 2 for trials in which the cued item was neutral, as a function of cue compatibility. Error bars denote within-subject standard errors (Morey, 2008).

##### Accuracy

Mirroring the RT data, the effect of the cued item was significant when the cue was neutral, F(1,13) = 15.28, p = 0.002, 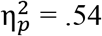. with conclusive evidence for this effect, *BF*_10_ = 39.61. The cue compatibility effect was also significant, F(2.26) = 5.32, p = .01, 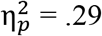 but the effect resulted from higher accuracy when the cue was neutral relative to when it was either compatible or incompatible, F(1, 13) = 4.94, p =.04, 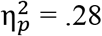 and F(1, 13) = 9.67, p =.008, 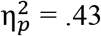, respectively. Crucially for the present purposes, there was no difference between the latter two conditions, F(1, 13) = 1.88, p =.19, 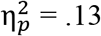, *BF*_10_ = 0.65.

#### Overall ANOVA

##### Reaction times

The main effect of cued item compatibility was significant, F(2, 26) = 140.78, p < .0001, 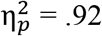: RTs were faster when the target rather than a neutral distractor appeared at the cue location, F(1, 13) = 101.76, p < .0001, 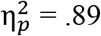 (cue-validity effect), and when a neutral rather than an incompatible distractor appeared there, F(1, 13) = 88.72, p < .0001, 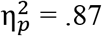 (cued distractor compatibility effect). The main effect of cue compatibility was not significant, F(2, 26) = 2.36, p = .11, 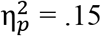. The interaction between the two factors was significant, F(4, 52) = 28.54, p < .0001, 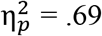.

Further analyses clarified this interaction. When the cued item was the target, the cue compatibility effect was significant, F(2, 26) = 29.46, p < .0001, 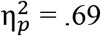. Paired comparisons revealed faster RTs when the cue was compatible than when it was incompatible, F(2, 26) = 51.4, p < .0001, 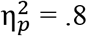. When the cued item was incompatible, the cue compatibility effect was also significant, F(2, 26) = 9.26, p = .001, 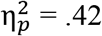, but the pattern of results was in the opposite direction: RTs were faster for incompatible than for compatible cues, F(1, 13) = 18.22, p = 001, 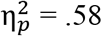.

##### Accuracy

The main effect of cued item was significant, F(2, 26) = 14.22, p <.0001, 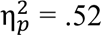, with a significant cued-distractor compatibility effect, F(1, 13) = 18.86, p < .001, 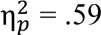, but no cue-validity effect, F < 1. The main effect of cue compatibility was also significant, F(2, 26) = 5.23, p = .01, 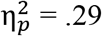: accuracy tended to be higher for compatible than for incompatible cues, F(1, 13) = 3.06, p = .08, 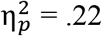, but was highest on neutral-cue trials. The interaction between the two factors was not significant F < 1.

### Discussion

The results of Experiment 2 closely replicated the results of Experiment 1. Thus, although we weakened the color and temporal clues associated with the search display, we failed to induce participants to rely on the detection of the target-defining color in order to deploy their attention. Participants still waited for the search display to appear, and must have relied on other differences between the cue and search display in order to discriminate the search-relevant context.

The findings nevertheless provide strong additional support for PAF. Indeed, whether we could interpret the results of Experiment 1 according to PAF critically depended on the specific pattern of cue-compatibility effects that is predicted by episodic retrieval accounts (e.g., Kahneman et al., 1992; Neill, 1997; van Dam & Hommel, 2010): a null cue-compatibility effect when a neutral distractor is cued, a positive effect (i.e., better performance on compatible than on incompatible cue trials) when the target is cued, and a negative effect (i.e., better performance on incompatible than on compatible cue trials) when an incompatible distractor is cued. Precisely this pattern was observed in Experiment 2.

Taken together, the findings of both experiments suggest that the deployment of spatial attention occurs in the search display, as predicted by PAF, and that the complex pattern cue-compatibility effects can be entirely accounted for by a backward process of episodic retrieval (e.g., Kahneman et al., 1992; Neill, 1997; van Dam & Hommel, 2010). Yet, further research is needed to examine whether contextual information triggers the deployment of attention, as is assumed by PAF.

## Experiment 3

The conclusion that the results of Experiments 1 and 2 support PAF rely on two findings. One is that the compatibility of the cue (in the cueing display) with the target did not affect performance - a finding that was clearest when response compatibility could be disentangled from episodic retrieval effects, that is, for neutral cued-distractor trials. The other is that the compatibility of the cued item strongly affected performance across conditions.

However, it was most probably easier to discriminate the direction of the large arrows (in the search display) that drove the cued item compatibility effects than the direction of the group of small arrows that made up the cue (in the cue display) and drove the cue compatibility effects. In other words, we may have failed to observe cue compatibility effects simply because the cue arrows were too difficult to discriminate.

The positive and negative cue compatibility effects observed when the target and an incompatible distractor, respectively appeared at the cue location seem to invalidate this alternative account: these effects attest to the fact that observers could discriminate the direction of the cue arrows. However, there is no principled reason to assume that the perceptual salience necessary to trigger a response dictated by the task at hand (forward effect) is the same as the perceptual salience necessary for memory traces to elicit match/mismatch effects when compared to the incoming input (backward episodic retrieval process).

It was therefore important to replicate our findings when the critical arrows in the cue display were highly salient. This was the purpose of Experiment 3. This experiment was similar to Experiment 1 except that the arrows in the cueing display were larger and thicker than the arrows in the search display. In addition, as in Experiment 2, we used multi-color cue displays.

### Methods

#### Participants

Fourteen new Tel Aviv University students (11 females, mean age = 22.71 years, SD = 2.67) participated in Experiment 3 as part of a course requirement, except for one participant who participated for payment. All participants were right-handed and reported normal or corrected-to-normal visual acuity.

#### Stimuli, procedure, design and statistical analysis

The sequence of events is presented in Figure 7. The stimuli, procedure and design were similar to those used for the regular trials of Experiment 2, with two exceptions: each arrow group of the cueing display was replaced with a single arrow-box (3.35°×3°) and the fixation placeholder was replaced with a gray fixation cross (0.25°×0.25°).

**Figure 7.**
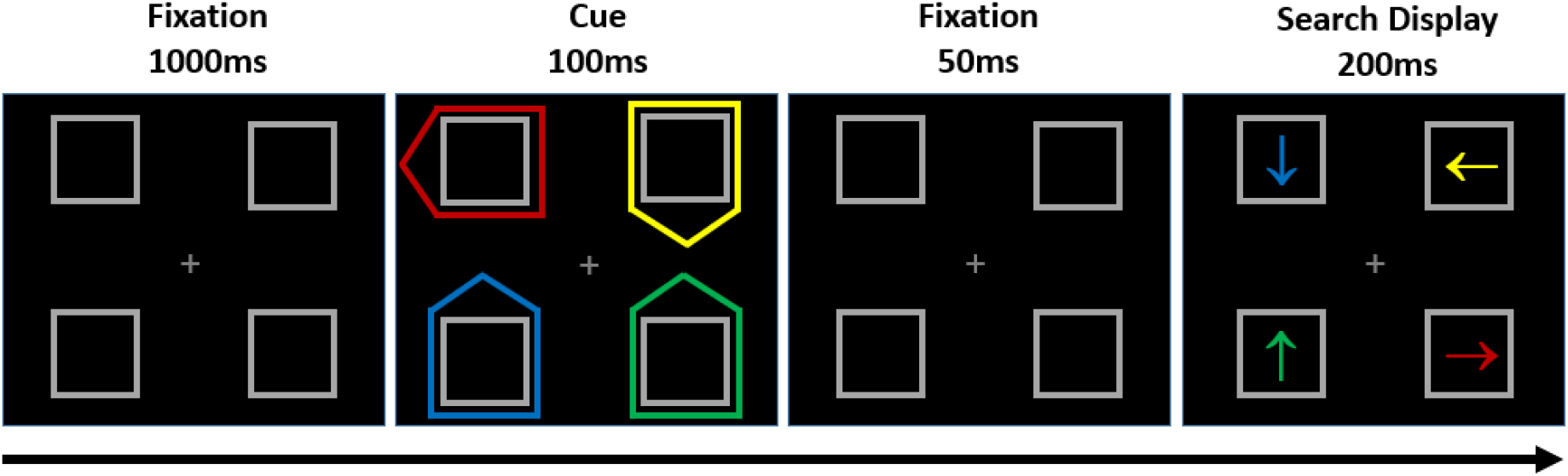
Sample sequence of events in Experiment 3, for the red-horizontal target group. Participants searched for the red arrow in the search display and reported its orientation (left or right). This example corresponds to an invalid-cue trial, in which the cue was incompatible with the target and the cued item was a neutral distractor.

### Results

No participant met the conditions for exclusion. Error trials (6.92% of the trials) as well as RT outliers (2.01 % of the remaining trials) were excluded from all RT analyses. Mean RTs and accuracy data are presented in Table 3.

**Table 3.**
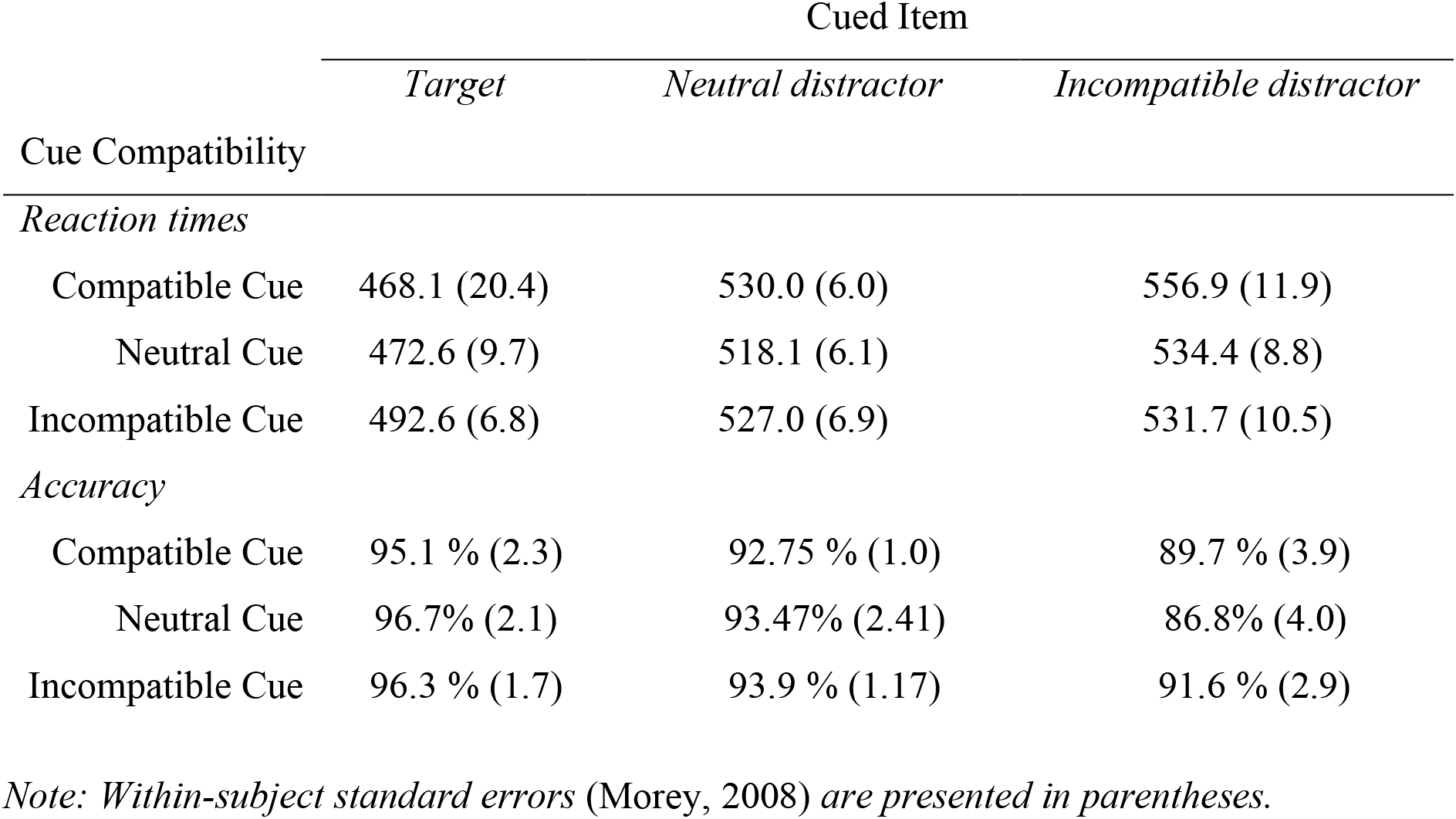
Mean reaction times (in milliseconds) and mean accuracy (in percentage) in Experiment 3, as a function of the compatibility of the cue with the target (Cue Compatibility: compatible, incompatible, neutral) and of the cued item in the search display (Cued Item: target, incompatible distractor, neutral distractor).

|  | Cued Item |  |  |
| --- | --- | --- | --- |
|  | *Target* | *Neutral distractor* | *Incompatible distractor* |
| Cue Compatibility |  |  |  |
| *Reaction times* |  |  |  |
| Compatible Cue | 468.1 (20.4) | 530.0 (6.0) | 556.9 (11.9) |
| Neutral Cue | 472.6 (9.7) | 518.1 (6.1) | 534.4 (8.8) |
| Incompatible Cue | 492.6 (6.8) | 527.0 (6.9) | 531.7 (10.5) |
| *Accuracy* |  |  |  |
| Compatible Cue | 95.1 % (2.3) | 92.75 % (1.0) | 89.7 % (3.9) |
| Neutral Cue | 96.7% (2.1) | 93.47% (2.41) | 86.8% (4.0) |
| Incompatible Cue | 96.3 % (1.7) | 93.9 % (1.17) | 91.6 % (2.9) |

*Note: Within-subject standard errors (Morey, 2008) are presented in parentheses.*

#### Planned comparisons

##### Reaction times

The results of Experiment 3 closely replicated the findings of Experiments 1 and 2. When the cue was neutral, the effect of cued distractor compatibility was highly significant, F(1, 13) = 15.12, p = 002, 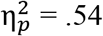, with conclusive evidence, *BF*_10_ = 15.96 (see Figure 8). When the cued item (in the search display) was a neutral distractor, the effect of cue compatibility was not significant, F < 1, with conclusive evidence for the null, *BF*_10_ = .326 (see Figure 9).

**Figure 8:**
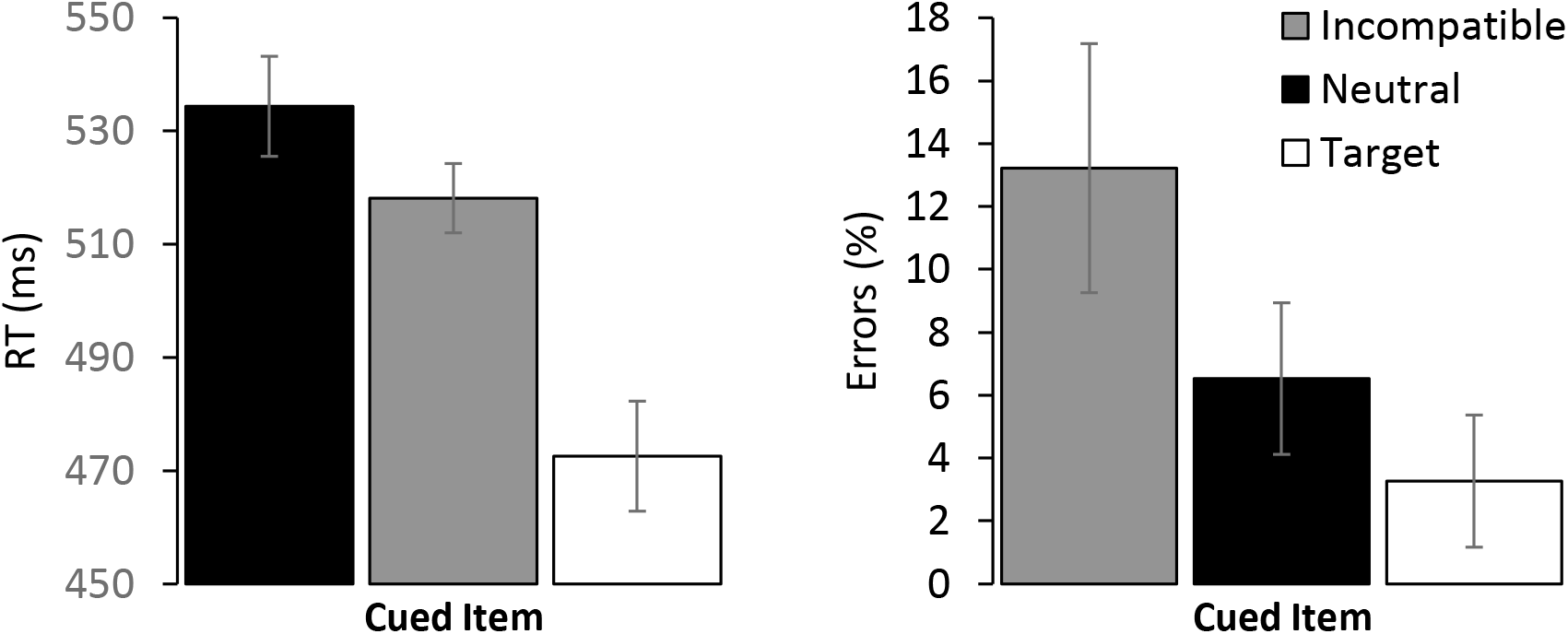
Mean reaction times (RT - left panel) and mean errors (right panel) in Experiment 3 for trials in which the cue compatibility was neutral, as a function of the cued item. Error bars denote within-subject standard errors (Morey, 2008).

**Figure 9:**
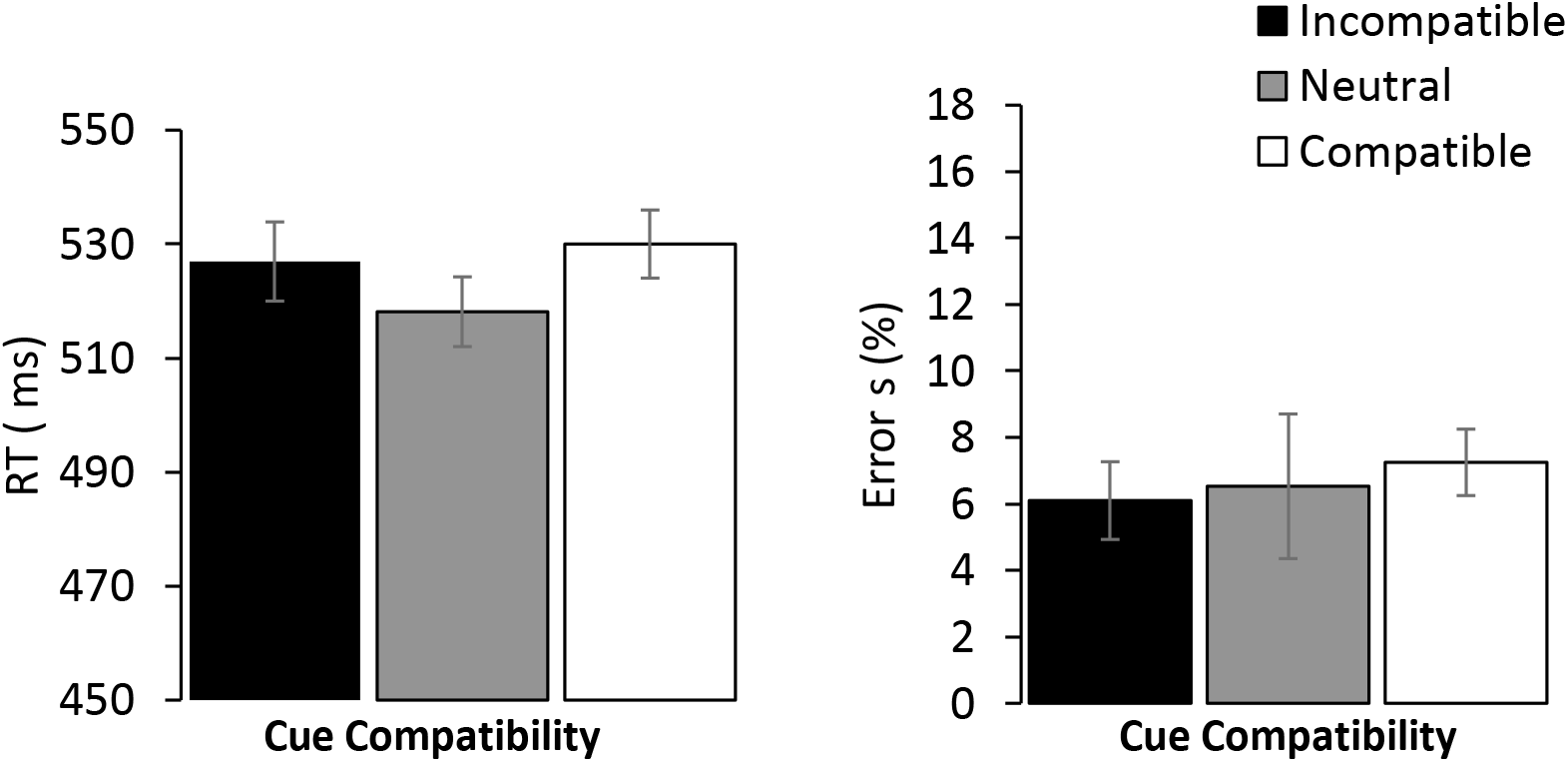
Mean reaction times (RT - left panel) and mean errors (right panel) in Experiment 3 for trials in which the cued item was neutral, as a function of cue compatibility. Error bars denote within-subject standard errors (Morey, 2008).

##### Accuracy

The accuracy data mirrored the RT data. When the cue was neutral, the effect of cued distractor compatibility was significant, F(1, 13) = 11.88, p = .004, 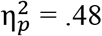, with conclusive evidence, *BF*_10_ = 10.7. When the cued item was a neutral distractor, the effect of cue compatibility was not significant, F < 1, yet the evidence for the null was inconclusive, *BF*_10_ = 1.09.

#### Overall ANOVA

##### Reaction times

The main effect of cued item was highly significant, F(2, 26) = 104.12, p < .0001, 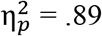: RTs were faster when the target rather than a neutral distractor appeared at the cued location, F(1, 13) = 98.13, p < .0001, 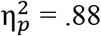 (cue-validity effect) and when a neutral rather than an incompatible distractor appeared there, F(1, 13) = 47.46, p < .0001, 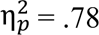 (cued-distractor compatibility effect). The main effect of cue compatibility was not significant, F(2, 26) = 3.11, p = .06, 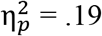. The interaction between cue compatibility and cued item was significant, F(4, 52) = 8.56, p < .0001, 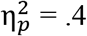.

As in Experiment 2, follow-up analyses revealed that when the cued item was the target, the cue compatibility effect was significant, F(2, 26) = 4.88, p = .016, 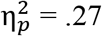, with slower RTs when the cue was incompatible than when it was compatible, F(1, 13) = 6.17, p = .027, 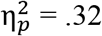. When the cued item was the incompatible distractor, there was again a significant negative cue compatibility effect, F(2, 26) = 8.66, p = .001, 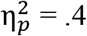 : RTs were faster than when the cue was incompatible than when it was compatible, F(1, 13) = 11.32, p = .005, 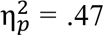.

##### Accuracy

The numerical trends on accuracy were similar to those observed on the RT data, thus removing any concern for a speed-accuracy trade-off. The main effect of cued item was significant, F(2, 26) = 21.56, p < .0001, 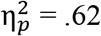, with significant effects of both its cue-validity and cued-distractor compatibility components, F(2, 26) = 8.31, p =.013, 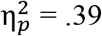 and F(2, 26) = 22.88, p < .001, 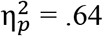, respectively. Neither the main effect of cue compatibility nor the interaction between the two factors was significant, F(2, 26) = 1.53, p = .24, 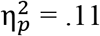 and F(4, 52) = 1.7, p = .63, 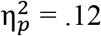, respectively.

### Discussion

In Experiment 3, the cue arrow was highly salient and easily discriminable. Yet, we replicated all the main findings of Experiment 1 and 2. When the two effects could not contaminate each other, the compatibility of the cued item affected responses to the target, whereas the compatibility of the cue did not. Moreover, the pattern of cue compatibility effects (positive when the target was cued and negative when the incompatible distractor was cued) again conformed to the episodic retrieval account (e.g., Kahneman et al., 1992). Thus, the results of Experiment 3 further invalidate the traditional interpretation of spatial cueing effects and support PAF’s claim that attention is deployed in the search display – not in the cue display.

### General Discussion

In three experiments, we presented evidence against the traditional interpretation of cue validity effects (e.g., Folk et al., 1992; Folk & Remington, 2006; Gaspelin et al., 2016; Theeuwes et al., 2000), according to which cue validity effects indicate that attention is allocated to the cue. We showed that attention is deployed only later, in the search display. These findings support a key hypothesis of the Priority Accumulation Framework (PAF, Gabbay et al., 2019; Lamy et al., 2018): as changes occur in the environment, we wait for relevant contextual information to deploy our attention.

#### Summary of the results

We used non-predictive cues sharing the target color in search for a target defined by its color and presented among heterogeneously colored distractors. Such cues are known to produce large and reliable validity effects (e.g., Carmel & Lamy, 2014; Folk & Remington, 1998; Lamy et al., 2004). We reasoned that if attention is allocated to the cue itself, the response-relevant feature of this cue should be processed and affect responses to the target. In other words, we should observe cue-target compatibility effects. By contrast, if attention is only deployed after the search display comes on, we should observe only effects of the compatibility between the cued item in the search display and the target (i.e., cued distractor compatibility effects). The results supported the latter prediction.

Specifically, the pattern of compatibility effects can be most parsimoniously explained as follows: the cue was not selected, as indicated by the null cue compatibility effect when the cued distractor was neutral (and could therefore not contaminate the effect of the cue). Instead, attention was deployed to the item that appeared at its location in the search display, as indicated by the cued-item compatibility effect that was significant across conditions of cue arrow directions. As a result, this item’s history was retrieved and response times depended on the match between its current and past response-relevant features (e.g., Gordon & Irwin, 1996.; Kahneman et al., 1992; Neill, 1997; Noles et al., 2005; van Dam & Hommel, 2010): when the target was cued, responses were faster if the cue was compatible (i.e., if it indicated the same direction as the target) than if it was incompatible, and conversely, when an incompatible distractor was cued, responses were faster if the cue was also incompatible (i.e., if both the cue and the distractor at its location indicated the same direction) than if it was compatible^2^.

#### Potential limitations

A first potential problem to note is that in all the present experiments, the cue display appeared for 100ms and was followed by a 50-ms blank before the search display appeared. Therefore, one may argue that a 150-ms SOA was not enough for response preparation to yield cue compatibility effects. However, this possibility is unlikely for two reasons. The first is that the cue arrows were not masked by the search display, therefore leaving ample time for them to take effect. The second reason is that the search display was also presented briefly, for 200ms, and yet, compatibility effects from the cued distractor were very robust. It is doubtful that a 50-ms difference in unmasked exposure times might explain why the cued item yielded reliable compatibility effects while the cue did not.

Second, one may also argue that response preparation to the arrow at the cued location overrode the compatibility effect generated by the cue. However, the null cue compatibility effect observed when the cued item was a neutral distractor argues against this conjecture, because neutral distractors were not associated with any response.

Third, Folk and Remington, (2006) reported findings that seem to contradict ours. They used a spatial cueing paradigm where the cue was a group of four small dots surrounding a placeholder box. On part of the trials that are of most interest here, the cue matched the target-defining color - as in the present study. Each search display contained two “X” and two “=” signs and participants had to report whether the color-defined target was an “X” or an “=”. Critically, on each trial, in the cueing display, a symbol (also either “X” or “=”) appeared within the placeholder that contained the cue (with all other placeholders remaining empty). This symbol could therefore be either compatible or incompatible with the target’s response feature. The authors found a significant cue compatibility effect on both valid- and invalid-cue trials. They concluded that attention was allocated to the cue in the cueing display. We replicated the cue compatibility effect on valid-cue trials. On invalid-cue trials, however, we found no cue compatibility effect when the cued distractor was neutral and a negative effect when this distractor was incompatible. By contrast, Folk and Remington, (2006) found a positive overall effect.

It should be noted that our findings and Folk and Remington’s (2006) are not necessarily inconsistent. In both studies, the cue compatibility effect was significant when the cue was valid. The discrepancy emerges when the cue was invalid. In our study, the distractor that appeared at the cue location (i.e., the cued distractor) could be either incompatible or neutral relative to the target (see footnote 1); we found a negative cue compatibility effect in the former condition and no effect in the latter. In Folk and Remington’s (2006) study, the cued distractor was either compatible or incompatible with the target but the authors did not report the cue compatibility effect separately for these two conditions. They only reported an overall positive cue compatibility effect on invalid-cue trials, which was smaller than on valid-cue trials and was not tested for significance. Thus, this small positive effect may reflect the net outcome of positive and negative effects (for compatible and incompatible cued distractor trials, respectively) – effects that we attributed to episodic retrieval processes occurring after attention is allocated to one of the search display’s objects (e.g., Kahneman et al., 1992). In addition, perceptual priming may have enhanced the cue compatibility effect in Folk and Remington’s study because unlike here, the symbols used in the cue and search displays were identical.

Finally, it should be noted that although we showed that attentional deployment occurs in the search display, we did not identify which contextual information associated with the search display triggered such deployment. Indeed, even though we made the temporal and display-wide characteristics of the search display less tractable (in Experiment 2), we could not trick our participants into allocating their attention in the cue display. However, other differences between the cue and search displays, for instance, the shape of the arrows, could help participants identify the search display. Further research should therefore provide stronger incentives for participants to mistakenly deploy their attention in the cue display. Nevertheless, our findings still confirmed our major hypothesis, namely, that attentional deployment occurred in the search display – and not in the cue display.

#### Relation to other models of attention

By showing that attentional deployment occurs in the search display rather than in the cue display, the present findings support PAF (Gabbay et al., 2019; Lamy et al., 2018) and challenge models that assume that at any given moment attention selects the object with the highest priority in the visual field (e.g., Folk & Remington, 2006; Gaspelin et al., 2016; Theeuwes, 2010; Treisman & Sato, 1990; Wolfe & Horowitz, 2017).

In recent series of papers, Zivony and Lamy (2016; 2018; Zivony et al., 2018) proposed a camera metaphor of attention that distinguishes between two successive operations during attentional allocation: shifting, during which attention is moved through space, and engagement, during which information in the focus of attention is processed. Most crucially for the present purposes, they suggested that the event that triggers attentional engagement is the detection of the target-defining feature. The evidence for this claim that is most relevant here was reported by Zivony and Lamy (2018). They used a spatial cuing paradigm, in which distractors were highly similar to the target - recent evidence shows that under such circumstances, an abruptly onset cue captures attention even if it does not share the target-defining feature (Gaspelin et al., 2016; Lamy et al., 2018). Zivony and Lamy observed both a cue validity effect and an effect of the compatibility between the cued distractor and the target when the onset cue shared the target-defining feature, but only a cue validity effect when the cue did not share this feature. They concluded that while goal-directed capture elicits both attentional shifts (indexed by cue validity effects) and attentional engagement (indexed by compatibility effects), stimulus-driven capture elicits “shallow shifts of attention”, that is, attentional shifts that are not followed by attentional engagement.

The present results, as well as previous findings supporting PAF (Lamy et al., 2018) call the camera metaphor into question on three accounts. First, we reported compatibility effects when the cue did not match the target-defining feature (Lamy et al., 2018). Second, here, we showed that detection of the target feature (in the cue display) does not trigger attentional engagement. Instead, attentional selection occurred in the search display, in line with PAF’s hypothesis that the search context signals the appropriate moment for deploying attention to the location that has accumulated the highest priority.

Finally, PAF suggests a more parsimonious account of the extant findings, as it does not require a distinction between attentional shifts and attentional engagement. In keeping with the core insight of the camera model (Zivony, Allon, Luria & Lamy, 2018; Zivony & Lamy, 2016), PAF also posits that cue validity and compatibility effects can be dissociated and are not two equivalent measures of attentional selection as is often assumed (e.g., Theeuwes et al., 2000). However, according to PAF, these measures do not reflect attentional shifting and engagement, respectively. Instead, cue validity effects indicate that the cue biases competition in the search display, whereas cued distractor compatibility effects indicate that attention is shifted to the location of the cued distractor and that, as a result, its features are processed. The PAF readily accounts Zivony and Lamy’s findings (2018): as target-matching cues are endowed with a larger priority weight than non-matching cues, a distractor at the location of a target-matching cue is most likely to win the competition (hence the observed cue validity and compatibility effects). In addition, target-distractor similarity was high enough for non-matching cues to bias the competition (hence, the observed spatial cueing effects), but too low for a distractor to win the competition when cued (hence, the null compatibility effects).

#### Concluding remarks

In this study we raised a hitherto neglected question in attention research: is spatial attention automatically moved to the ever-changing location with the highest priority at any given moment? Or does its deployment await a trigger that signals the appropriate context? Our results suggest that the latter is true. As such, they dovetail findings from temporal attention research (e.g., Coull & Nobre, 1998) suggesting that we can withhold the allocation of attention until the most appropriate moment has arrived for selection to best serve our search goals. Our findings call for a revision of leading models of attention that overlook the role of search context in the timing of attentional allocation. Although some of these models do posit a role for contextual information (e.g., Wolfe & Horowitz, 2017), such information is thought to determine what should be selected rather than when.

## Author’s note

Correspondence should be addressed to Dominique Lamy at. Support was provided by the Israel Science Foundation (ISF) grants no. 1286/16 to Dominique Lamy.

## Footnotes

1. There was no compatible distractor because there was only one arrow in each direction in the search display. Adding a compatible distractor would open the way to an alternative strategy for responding correctly: the arrow direction of which there are two exemplars would always be the target-arrow direction.

2. 2. An auxiliary finding worth noting is that a cue sharing the target-defining feature produced substantial validity effects even when it was not salient, that is, when it appeared among heterogeneously colored objects and was therefore not a singleton (55ms in Exp.2 and 46ms in Exp.3). This finding (also reported by Lamy et al., 2004) demonstrates the powerful role of top-down factors in guiding attention and therefore provides strong support for contingent capture (e.g., Folk et al., 1992).

## References

Carmel, T., & Lamy, D. (2014). The same-location cost is unrelated to attentional settings: An object-updating account. Journal of Experimental Psychology: Human Perception and Performance, 40(4), 1465–1478. https://doi.org/10.1037/a0036383

Cave, K. R., & Wolfe, J. M. (1990). Modeling the role of parallel processing in visual search. Cognitive Psychology, 22(2), 225–271. https://doi.org/10.1016/0010-0285(90)90017-X

Coull, J. T., & Nobre, A. C. (1998). Where and when to pay attention: The neural systems for directing attention to spatial locations and to time intervals as revealed by both PET and fMRI. Journal of Neuroscience, 18(18), 7426–7435. https://doi.org/10.1523/jneurosci.18-18-07426.1998

Dienes, Z., & Mclatchie, N. (2018). Four reasons to prefer Bayesian analyses over significance testing. Psychonomic Bulletin and Review, 25(1), 207–218. https://doi.org/10.3758/s13423-017-1266-z

Faul, F., E. Erdfelder, A. Buchner, and A. G. Lang. 2013. “G* Power Version 3.1. 7 [Computer Software].” Uiversität Kiel, Germany.

Folk, C. L., & Remington, R. (1998). Selectivity in Distraction by Irrelevant Featural Singletons: Evidence for Two Forms of Attentional Capture. Journal of Experimental Psychology: Human Perception and Performance, 24(3), 847–858. https://doi.org/10.1037/0096-1523.24.3.847

Folk, C. L., & Remington, R. (2006). Top-down modulation of preattentive processing: Testing the recovery account of contingent capture. In Visual Cognition (Vol. 14, Issues 4–8). https://doi.org/10.1080/13506280500193545

Folk, C. L., Remington, R. w, & Johnston, J. C. (1992). 1992_Folk_etal_JEPHPP.pdf. In Journal of Experimental Psychology: Human Perception and Performance (Vol. 18, Issue 4, pp. 1030–1044).

Gabbay, C., Zivony, A., & Lamy, D. (2019). Splitting the attentional spotlight? Evidence from attentional capture by successive events*. Visual Cognition, 27(5–8), 518–536. https://doi.org/10.1080/13506285.2019.1617377

Gaspelin, N., Ruthruff, E., & Lien, M.-C. (2016). The problem of latent attentional capture: Easy visual search conceals capture by task-irrelevant abrupt onsets. In Journal of Experimental Psychology: Human Perception and Performance (Vol. 42, Issue 8, pp. 1104–1120). American Psychological Association. https://doi.org/10.1037/xhp0000214

Gibson, B. S., & Kelsey, E. M. (1998). Stimulus-Driven Attentional Capture Is Contingent on Attentional Set for Displaywide Visual Features. Journal of Experimental Psychology: Human Perception and Performance, 24(3), 699–706. https://doi.org/10.1037/0096-1523.24.3.699

Gordon, R. D., & Irwin, D. E. (1996). What’s in an object file? Evidence from priming studies. Perception and Psychophysics, 58(8), 1260–1277. https://doi.org/10.3758/BF03207558

Green, E. J., & Quilty-Dunn, J. (2017). WHAT IS AN OBJECT FILE? http://www.nyu.edu/gsas/dept/philo/courses/readings/2016.green.qd.pdf

Hommel, B. (1998). Event files: Evidence for automatic integration of stimulus-response episodes. Visual Cognition, 5(1–2), 183–216. https://doi.org/10.1080/713756773

Itti, L., & Koch, C. (2000). A saliency-based search mechanism for overt and covert shifts of visual attention. Vision Research, 40(10–12), 1489–1506. https://doi.org/10.1016/S0042-6989(99)00163-7

Kahneman, D., & Henik, A. (2017). Perceptual organization and attention. In Perceptual organization (pp. 181–211). Routledge.

Kahneman, D., Treisman, A., & Gibbs, B. J. (1992). The reviewing of object files: Object-specific integration of information. Cognitive Psychology, 24(2), 175–219. https://doi.org/10.1016/0010-0285(92)90007-O

Lamy, D. (2005). Temporal expectations modulate attentional capture. Psychonomic Bulletin and Review, 12(6), 1112–1119. https://doi.org/10.3758/BF03206452

Lamy, D., Darnell, M., Levi, A., & Bublil, C. (2018). Testing the attentional dwelling hypothesis of attentional capture. Journal of Cognition, 1(1).

Lamy, D., Leber, A., & Egeth, H. E. (2004). Effects of task relevance and stimulus-driven salience in feature-search mode. Journal of Experimental Psychology: Human Perception and Performance, 30(6), 1019–1031. https://doi.org/10.1037/0096-1523.30.6.1019

Lamy, D, Leber, A., & Egeth. H.E. (2012). “Selective Attention.” Handbook of Psychology, Second Edition 4.

Morey, R. D. (2008). Confidence Intervals from Normalized Data: A correction to Cousineau (2005). Tutorials in Quantitative Methods for Psychology, 4(2), 61–64. https://doi.org/10.20982/tqmp.04.2.p061

Morey, R. D., Rouder, J. N., Jamil, T., & Morey, M. R. D. (2015). Package ‘bayesfactor.’ URLh Http://Cran/r-Projectorg/Web/Packages/BayesFactor/BayesFactor Pdf i (Accessed 1006 15).

Neill, W. T. (1997). Episodic retrieval in negative priming and repetition priming. Journal of Experimental Psychology: Learning, Memory, and Cognition, 23(6), 1291–3105. https://doi.org/10.1037/0278-7393.23.6.1291

Neisser, U. (1976). Cognition and reality: Principles and implications of cognitive psychology. In Cognition and reality: Principles and implications of cognitive psychology. W H Freeman/Times Books/ Henry Holt & Co.

Noles, N. S., Scholl, B. J., & Mitroff, S. R. (2005). The persistence of object file representations. Perception & Psychophysics, 67(2), 324–334.

Olson, I. R., & Chun, M. M. (2001). Temporal Contextual Cuing of Visual Attention. Journal of Experimental Psychology: Learning Memory and Cognition, 27(5), 1299–1313. https://doi.org/10.1037/0278-7393.27.5.1299

Theeuwes, J. (2010). Top-down and bottom-up control of visual selection. Acta Psychologica, 135(2), 77–99. https://doi.org/10.1016/j.actpsy.2010.02.006

Theeuwes, J., Atchley, P., & Kramer, A. F. (2000). On the time course of top-down and bottom-up control of visual attention. In Attention and Performance (Vol. 18, Issue February 2014).

Treisman, A. M., & Gelade, G. (1980). A feature-integration theory of attention. Cognitive Psychology, 12(1), 97–136.

Treisman, A., & Sato, S. (1990). Conjunction Search Revisited. Journal of Experimental Psychology: Human Perception and Performance, 16(3), 459–478. https://doi.org/10.1037/0096-1523.16.3.459

van Dam, W. O., & Hommel, B. (2010). How object-specific are object files? Evidence for integration by location. Journal of Experimental Psychology: Human Perception and Performance, 36(5), 1184–1192. https://doi.org/10.1037/a0019955

Wolfe, J. M., & Horowitz, T. S. (2017). Five factors that guide attention in visual search. Nature Human Behaviour, 1(3), 1–8. https://doi.org/10.1038/s41562-017-0058

Yantis, S., & Egeth, H. E. (1999). On the distinction between visual salience and stimulus-driven attentional capture. Journal of Experimental Psychology: Human Perception and Performance, 25(3), 661–676. https://doi.org/10.1037/0096-1523.25.3.661

Zivony, A., Allon, A. S., Luria, R., & Lamy, D. (2018). Dissociating between the N2pc and attentional shifting: An attentional blink study. Neuropsychologia, 121(August), 153–163. https://doi.org/10.1016/j.neuropsychologia.2018.11.003

Zivony, A., & Lamy, D. (2016). Attentional capture and engagement during the attentional blink: A “camera” metaphor of attention. Journal of Experimental Psychology: Human Perception and Performance, 42(11), 1886–1902. https://doi.org/10.1037/xhp0000286

Zivony, A., & Lamy, D. (2018). Contingent Attentional Engagement: Stimulus- and Goal-Driven Capture Have Qualitatively Different Consequences. Psychological Science, 29(12), 1930–1941. https://doi.org/10.1177/0956797618799302

